# His bundle has shorter chronaxie than adjacent ventricular myocardium: Implications for pacemaker programming

**DOI:** 10.1101/623348

**Authors:** Marek Jastrzębski, Paweł Moskal, Agnieszka Bednarek, Grzegorz Kiełbasa, Pugazhendhi Vijayaraman, Danuta Czarnecka

## Abstract

**Background:** Strength-duration curves for permanent His bundle (HB) pacing are potentially important for pacemaker programming.

**Objective:** We aimed to calculate strength-duration curve and chronaxie of the His bundle (HB) and of the adjacent right ventricular (RV) working myocardium and to analyze zones of selective HB capture and battery current drain when pacing at different pulse durations (PDs).

**Methods:** Consecutive patients with permanent HB pacing were studied. The RV and HB capture thresholds were assessed at several PDs. Battery current drain and zones of selective HB capture at PDs of 0.1, 0.2, 0.4 and 1.0 ms were determined.

**Results:** In the whole group (n =127) the HB chronaxie was shorter than the RV chronaxie. This difference was driven by patients with selective HB pacing (HB chronaxie of 0.47 vs RV chronaxie of 0.79 ms). Strength-duration curve for HB had lower rheobase and its steep portion started at shorter PDs thus creating wider distance - zone of programmable selective HB pacing - between the HB and RV strength-duration curves at shorter PDs. The battery current drain was lower with pacing at PDs of 0.1 - 0.4 ms vs 1.0 ms. Chronaxie adjusted PDs offered lowest current drain.

**Conclusion:** For the first time the strength-duration curves for permanent selective and non-selective HB pacing were determined. Selective HB capture and battery longevity can be promoted by shorter PDs (0.2 ms). Longer PDs (1.0 ms) offer bigger safety margin for RV capture and may be preferable if simultaneous RV capture during HB pacing is desired.

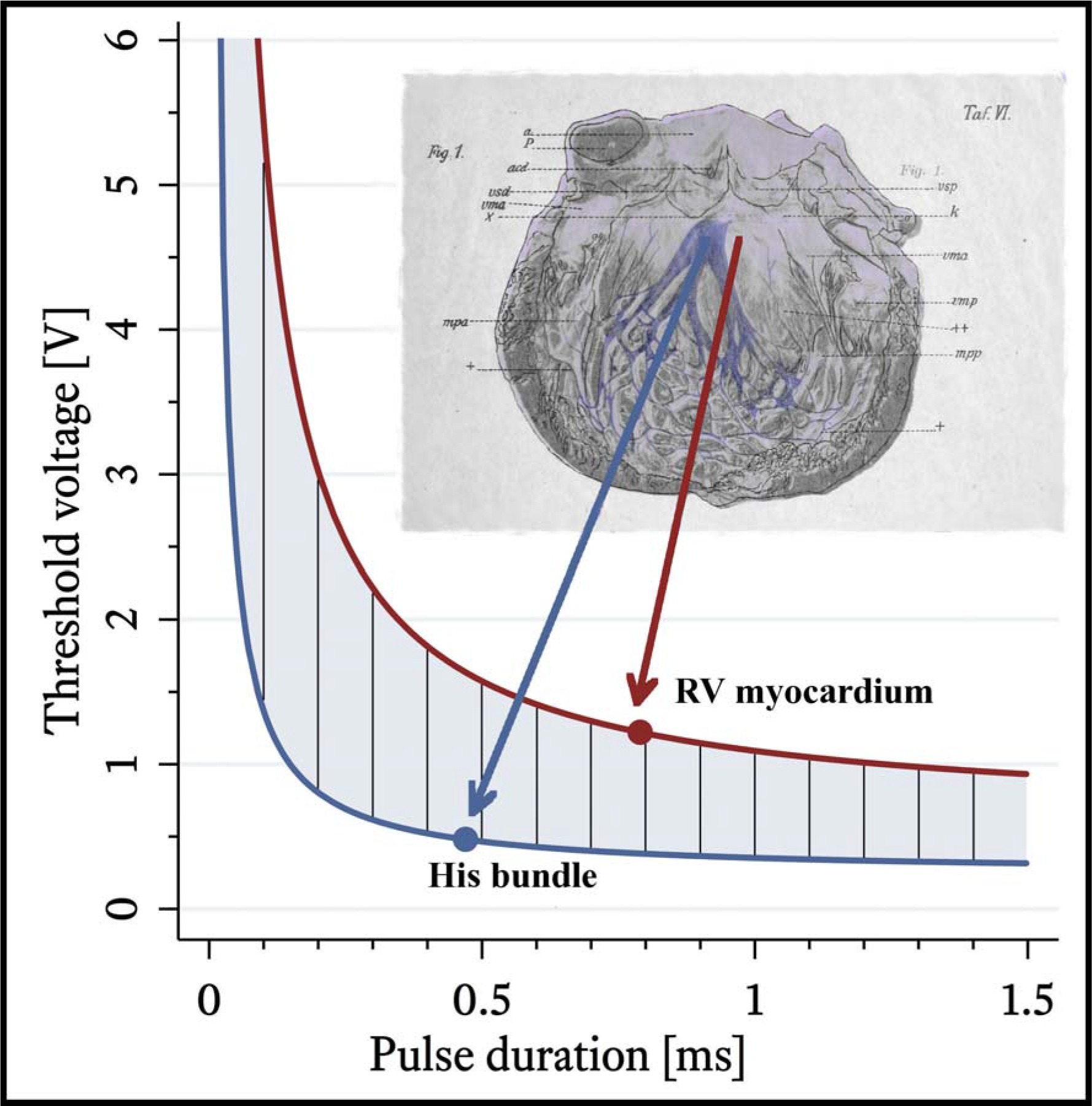

## Introduction

Permanent His bundle (HB) pacing offers most physiological spread of depolarization during ventricular pacing. It is free from many of the known detrimental effects of conventional right ventricular (RV) pacing, and might reverse the clinical course and structural changes induced by RV pacing and bundle branch blocks ^1–5^ With the growing amount of patients with permanent HB pacing devices ^6–11^ determination of basics of this form of pacing like chronaxie, rheobase and strength-duration curves, seems very important. Rheobase is defined as the smallest voltage amplitude that stimulates the tissue at an infinitely long pulse duration. Chronaxie is a tissue-specific measure that is used to describe the relative excitability of tissues. It is defined as the pulse duration at which the current required for stimulation is twice the rheobase. HB conduction is several times faster than working myocardium - and is known that chronaxie is shorter for rapidly propagating tissues. Pacing with pulse duration close to chronaxie is most efficient with regard to optimizing battery longevity. Following this rule might be especially important during HB pacing as high pacing thresholds and faster battery depletion is seen as major drawbacks of this pacing modality. However, to our knowledge, no study has reported chronaxie values during permanent HB pacing.

Another area where knowledge of strength-duration curves for HB might play a role is the achievement of selective vs. non-selective HB capture. A few studies reported that differences in chronaxie values enable selective pacing of different tissues from the same electrode by using a different combination of pulse widths and voltage output; whether this is also the case with His bundle/RV myocardium capture is not known.

We hypothesized that chronaxie is shorter for the HB than for the adjacent septal RV myocardium, and that pacing at pulse duration closer to chronaxie would both decrease battery current drain and facilitate selective HB capture. Our primary aim was to calculate strength-duration curves and determine chronaxies of the HB and of the adjacent RV myocardium during permanent HB pacing. Our secondary aim was to compare battery current drain and ranges of programmable zones of selective HB capture when pacing at pulse durations shorter and longer than chronaxie.

## Methods

The study was approved by the Institutional Bioethical Committee. Consecutive patients who underwent permanent HB pacing device implantation in our institution in the years 2014-2018 were screened and invited to participate in the study. In all cases Medtronic (MN, USA) SelectSecure 3830 lead was used for HB pacing. This is a 4.1F, active helix, steroid eluting lead with the cathode area of 3.6 mm.^2^ HB device implantation was performed using acknowledged methodology.^12, 13^ Basic clinical data including age, gender, co-morbidities, pacing indication, pacing mode and echocardiographic data were obtained from our prospectively maintained HB pacing database.

The first step of the study was determination of several points (minimum four) on the strength-duration curves, separately for HB pacing and for RV pacing. For this, both the RV and HB capture thresholds were assessed throughout the whole programmable voltage range at several pulse widths during VVI unipolar pacing. Threshold assessment in each patient was performed according to uniform follow-up charts (Figure 1) that included measurements at the following pulse widths: 1.5 ms, 1.0 ms, 0.4 ms, 0.3 ms, 0.2 ms, 0.1 ms or the closest values available in a particular device. Additional points (@ 0.5 ms, @ 0.75 ms) were assessed when pacing at shorter pulse duration was not available in the device program or there was no capture at short pulse durations.

**Figure 1.**
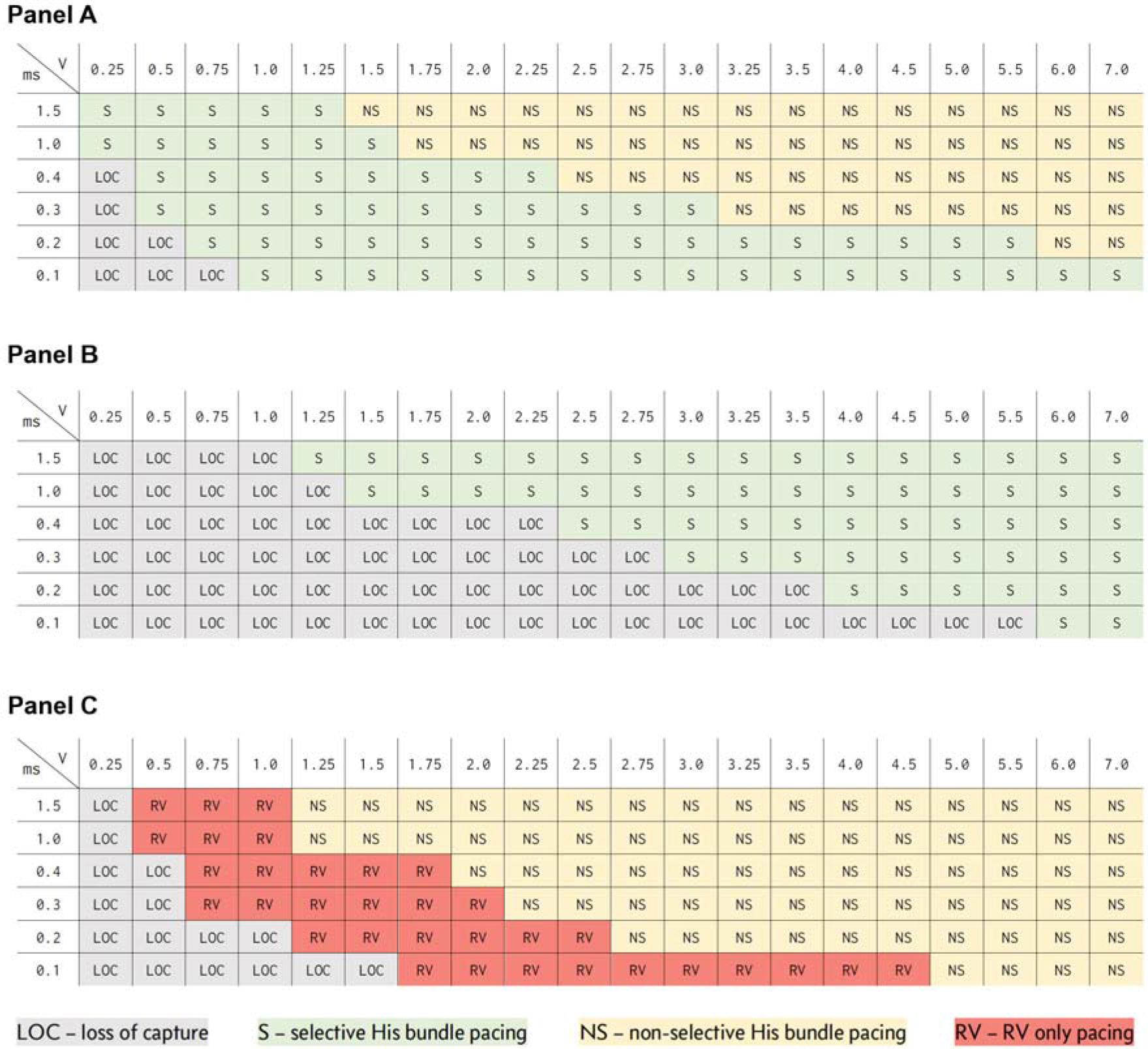
Threshold assessment charts for patients with permanent HB pacing illustrating three distinct patterns of capture with regard to selective / non-selective capture. Panel A: patient with wide zone of selective pacing, that broadens with shorter pulse width. Panel B: patient with obligatory selective capture. Panel C: patient with obligatory non-selective HB capture.

Simultaneous recording of all 12 standard ECG leads and observation of QRS morphology change supplemented by near-field EGM recording from pacemaker programmer were used as the primary method for differentiation between pure RV capture, selective HB capture and non-selective HB capture.^14, 15^ Programmed His bundle pacing method for differentiation between RV QRS and non-selective HB QRS was applied when considered necessary.^16^ Selective and non-selective HB capture was identified according to the recently proposed criteria.^17^ For the purpose of this study, a patient was categorized as having selective HB capture lead position if HB threshold was below RV threshold with at least one voltage and pulse width combination. The RV capture threshold was considered to be reached at a point of the transition from RV capture to loss of capture (RV → LOC) or at a point of sudden QRS narrowing (and appearance of isoelectric line after pacing stimulus) with pacing current decrease (non-selective HB → selective HB). The HB capture threshold was established by observation of transition from selective HB capture to loss of capture (HB → LOC) or at a point of QRS widening with pacing current increase (non-selective HB → RV).

For threshold measurement protocol, we followed “Ten Rules for Good Measuring Practice” formulated by W. Irnich,^18^ with the main following points:

1. smallest possible voltage steps, as allowed by the implanted device, were used to determine the capture threshold for RV and HB.
2. to increase precision, the threshold amplitude was recorded as an arithmetically averaged value of the last effective and the first ineffective voltage setting^18–20^
3. voltage thresholds were expressed as the mean voltage over pulse duration (called *quantity)*, where mean voltage value was estimated using the formula:

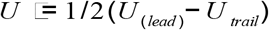

with:

U_lead_: leading edge voltage amplitude, set in pacemaker output setting
U_trail_: trailing edge voltage amplitude estimated from output waveform at nominal conditions with the assumption of linear voltage decline ^18–20^

In the second step of the study, the established RV and HB capture thresholds at different pulse widths served for:

1. determination of chronaxie (t_c_) and rheobase 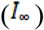 using linear regression calculation of the *quantity* versus pulse duration ^18–21^

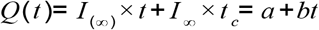

where:

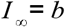

1. calculation of the strength-duration curves. i.e. plots of the threshold current versus pulse duration required to stimulate RV and HB
2. determination of the zones of selective HB capture at different pulse widths. Zone of selective HB capture, expressed in Volts, was defined as the range of pulse amplitudes that resulted in selective HB capture. Such zone starts with HB threshold amplitude and ends with RV threshold amplitude at a particular pulse width.
3. determination of the percentage of patients who would have selective HB capture when pacing at different pulse widths when following the rules of pacing with safety margin defined as pacing output set at HB capture threshold + 1 V or as twice the HB capture threshold (with minimal voltage output at 1 V).
4. calculation of current drain when pacing at different pulse widths following the rules of pacing with a safety margin. Battery current drain - I_dr_ (as measured in microampere-hours, μAh) by a single pacing stimulus was calculated according to the formula:

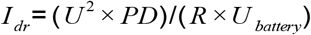

Where: U is output voltage (V), PD is the pulse duration (ms), R is output impedance 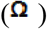 and U_battery_ is a uniform battery voltage value of 2.8 V.

### Statistical methods

Basic and clinical data are reported as mean ± SD in case of normal distribution, which was assessed by Shapiro–Wilk test. Chronaxie and rheobase are presented as median ± interquartile range. Wilcoxon matched-pairs signed-ranks test was used to compare chronaxie, rheobase, programmable zones of selective His bundle pacing width and mean pulse current drain. *P* values of < 0.05 were considered statistically significant.

## Results

One hundred twenty-seven patients who underwent successful implantation of HB pacing device were studied. These patients were mostly older, with many co-morbidities and moderately depressed left ventricular ejection fraction, almost all had narrow QRS complexes. HB pacing devices were implanted mainly for permanent atrial fibrillation with bradycardia or proximal atrioventricular block with narrow QRS complexes. Basic clinical and procedure related data are presented in greater detail in Table 1.

**Table 1.** Basic clinical and pacing related characteristics of the studied group.

| Characteristic | n = 127 |
| --- | --- |
| Age [years] | 73.7 ± 11.4 |
| Male gender | 91 (71.6%) |
| Heart failure | 66 (52.0%) |
| Hypertension | 107 (84.2%) |
| Coronary heart disease | 54 (42.5%) |
| Diabetes mellitus | 46 (36.2%) |
| Pacing indications: |  |
| • Atrial fibrillation with bradycardia | 65 (51.2%) |
| • Atrioventricular block | 25 (19.7%) |
| • Heart failure | 25 (19.7%) |
| • Sick sinus syndrome | 12 (9.4%) |
| EF [%] | 46.6 ± 14.0 |
| LVEDD [mm] | 54.6 (9.3%) |
| Preimplantation QRS duration width [ms] | 112.4 ± 25.5 |
| Paced QRS complex width [ms] | 119.5 ± 23.1 |
| HB capture threshold @0.4 ms pulse duration [V] | 1.16 ± 0.83 |
| Length of follow-up [months] | 17.9 ± 15.8 |

EF - ejection fraction; LVEDD – left ventricle end-diastolic dimension; AF – atrial fibrillation;

### Chronaxie and rheobase

The results of chronaxie and rheobase measurements in the whole group and in the subset of selective HB pacing and obligatory non-selective HB pacing are presented in Table 2.

**Table 2.** His bundle and right ventricular chronaxie and rheobase values in subgroups

|  | RV capture | HB capture | P value |
| --- | --- | --- | --- |
| **Chronaxie [ms]** |  |  |  |
| whole group | 0.77 (0.58–1.15) | 0.53 (0.32–0.76) | 0.00001 |
| selective HB pacing | 0.79 (0.60–1.16) | 0.47 (0.28–0.71) | 0.00001 |
| obligatory non-selective HB pacing | 0.70 (0.46–1.03) | 0.72 (0.51–0.88) | ns |
| **Rheobase [V]** |  |  |  |
| whole group | 0.49 (0.21–0.90) | 0.26 (0.18–0.52) | 0.0001 |
| selective HB pacing | 0.61 (0.32–1.13) | 0.24 (0.17–0.44) | 0.0001 |
| obligatory non-selective HB pacing | 0.19 (0.17–0.24) | 0.45 (0.31–0.71) | 0.0001 |

RV – right ventricle, HB – His bundle, ns – non-significant

Selective HB pacing was seen in 99 (78%) patients, obligatory non-selective HB pacing (RV threshold always below HB threshold at all voltage and pulse width combinations) in 28 (21%). Strength-duration curves for RV and HB differed, as presented in Figure 2 and in Table 2.

**Figure 2.**
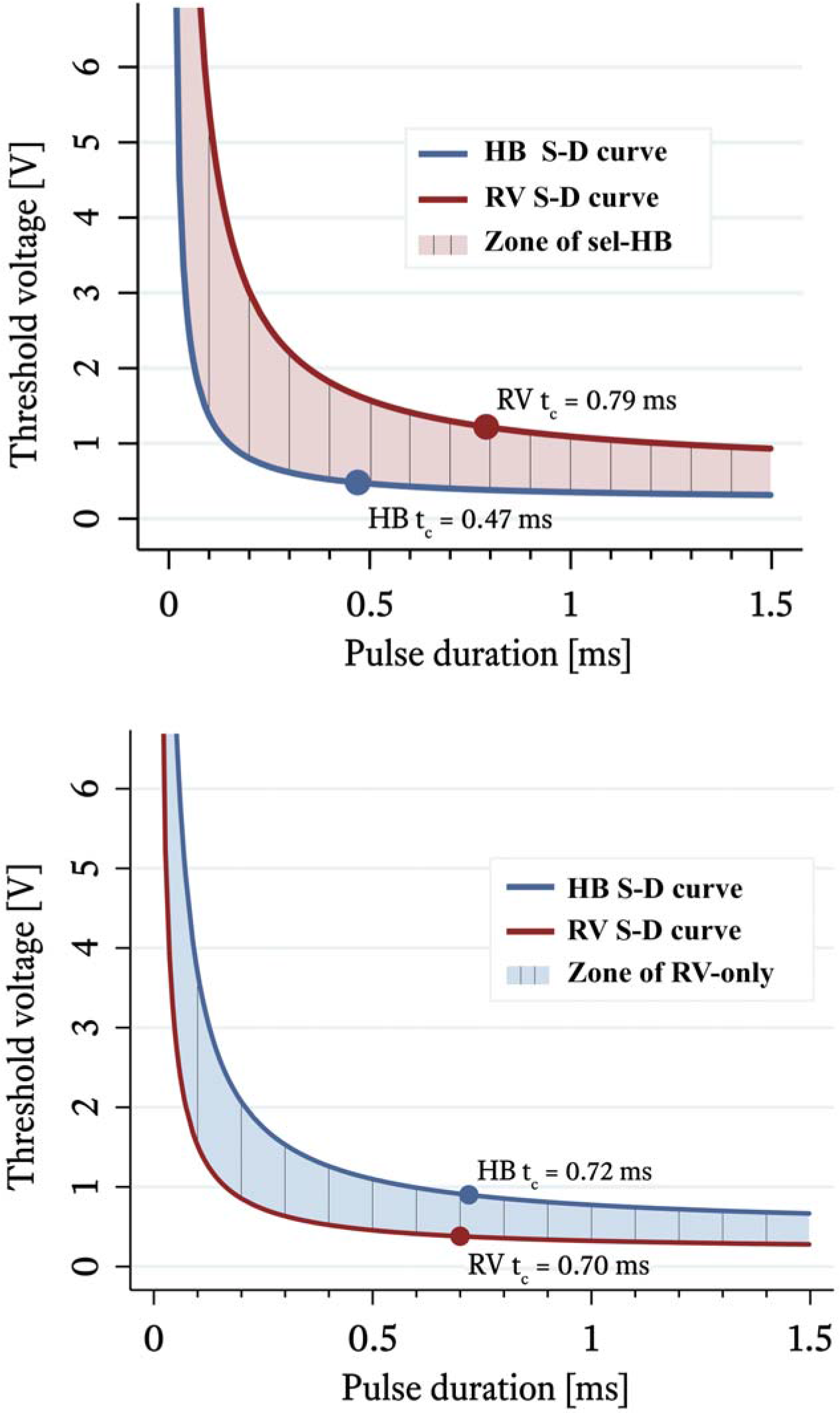
Upper panel: in patients with selective His bundle (HB) pacing chronaxie for HB is shorter than for right ventricular myocardium (RV). Consequently, zone of selective pacing widens with shortening of the pulse duration. Lower panel: in patients with obligatory non-selective HB pacing chronaxie values between HB and RV do not differ and zone of RV-only capture is more constant. S-D – strength-duration, T_c_ – chronaxie

The hyperbolic curve for HB had lower rheobase and its steep portion started at shorter pulse widths creating wider distance (zone of selective capture) between the curves at shorter pulse widths. HB chronaxie was shorter when compared to RV chronaxie, 0.53 ms vs 0.77 ms in the whole group. This was driven by the shorter HB chronaxie in patients with selective pacing (0.47 vs 0.79 ms, respectively) as chronaxie in patients with obligatory non-selective capture did not differ (0.72 vs 0.70 ms). Patients with selective HB capture had lower HB chronaxie than patients with obligatory non-selective HB capture (0.47 vs 0.72 ms), while there was no difference in RV chronaxie between these two subgroups. The calculated chronaxie range in patients with selective HB capture was 0.10 – 1.47 ms and 0.24 – 1.50 ms in patients with obligatory non-selective HB capture.

In the whole group, the RV rheobase was higher than the HB rheobase value (0.49 vs 0.26 V), however, in the subgroup with obligatory non-selective pacing, a reversed situation was observed (0.19 vs. 0.45 V). The HB rheobase was lower in selective HB pacing (0.24 vs 0.45 V, p = 0.006) and the RV rheobase was lower in obligatory non-selective HB vs selective HB (0.61 vs 0.19 V, p = 0.0001) - Table 2.

### Programming for selective His bundle pacing

Zone of programmable selective HB pacing was wider when pacing at shorter pulse duration (0.1 vs 0.2 ms, 0.2 vs 0.4 ms, 0.4 vs 1.0 ms); this data is pertinent to the selective HB pacing subgroup only - Figure 3.

**Figure 3.**
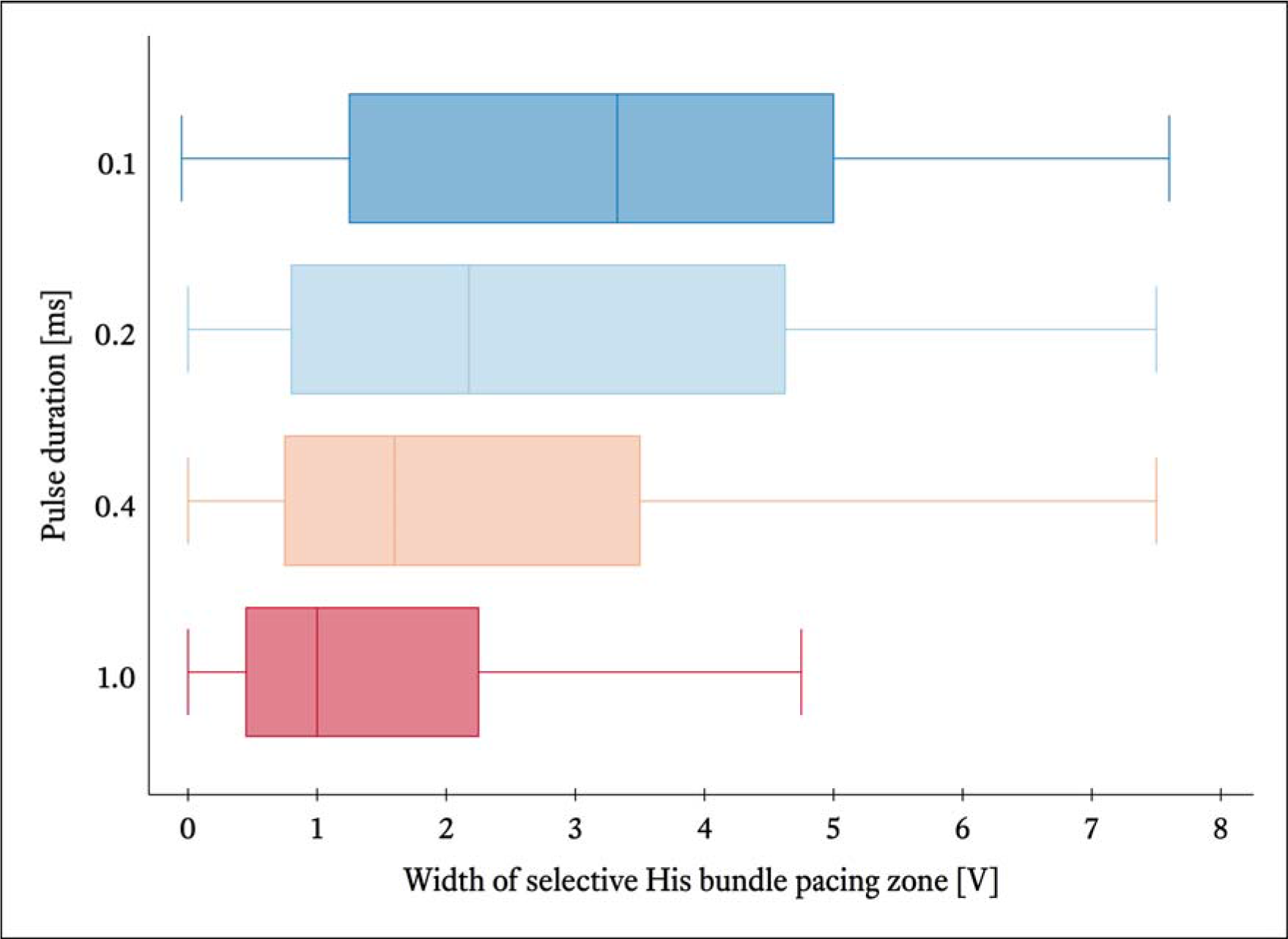
Programmable zones of selective His bundle capture pacing are wider for shorter pulse widths. Box plot with median and inter-quartile range (25% - 75%).

The percentage of patients with selective His bundle pacing during follow-up depends on pulse duration and safety margin programming. More patients had selective HB capture when paced at pulse duration of 0.1 ms and 0.2 than at 0.4 or 1.0 ms - when output was programmed with 1 V of safety margin (Figure 4, left panel). When the output voltage was set to twice the HB capture threshold voltage, the highest percentage of selective His bundle pacing cases where at PD of 0.4 ms (Figure 4, right panel).

**Figure 4.**
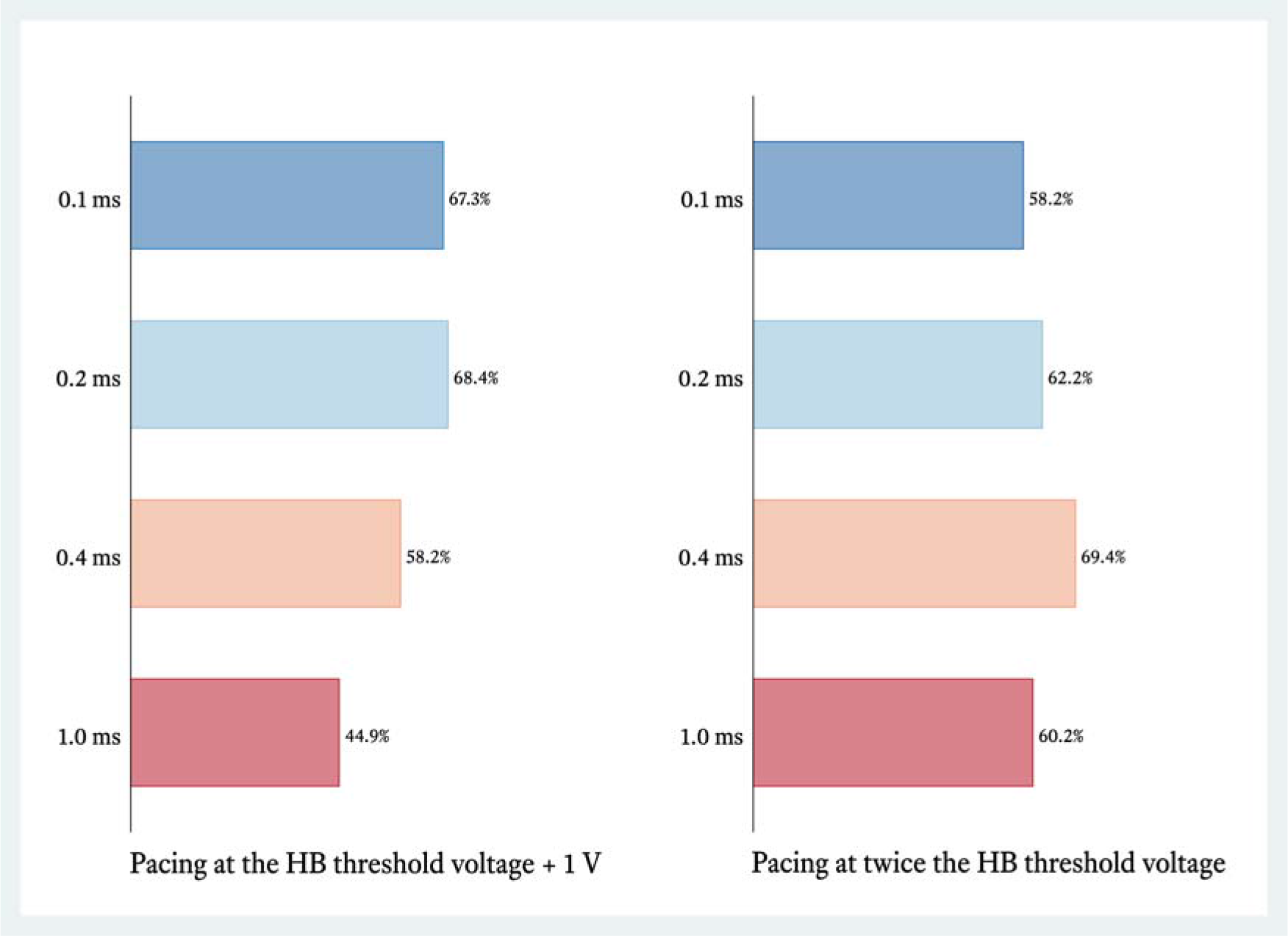
Percentage of patients with selective His bundle capture is related to programmed pulse widths and safety margin.

### Battery current drain

The battery current drain was lower (p = 0.0002) when pacing was programmed with a pulse duration of 0.1, 0.2, 0.4 ms vs 1.0 ms either following the rule of threshold +1 V (p = 0.0001) or twice the voltage threshold - Figure 5. However, the lowest current drain of 1.4 (0.44-3.48) uAh/pulse was obtained for safety rule of + 1 V when the pulse duration was individually selected for each patient according to calculated chronaxie (Table 3).

**Table 3.** The algorithm for adjusting pulse duration programming according to His bundle chronaxie

| His bundle chronaxie | Optimized pulse duration |
| --- | --- |
| < 0.4 ms | 0.2 ms |
| (0.4 ms - 1.0 ms) | 0.4 ms |
| ≥ 1.0 ms | 1.0 ms |

**Figure 5.**
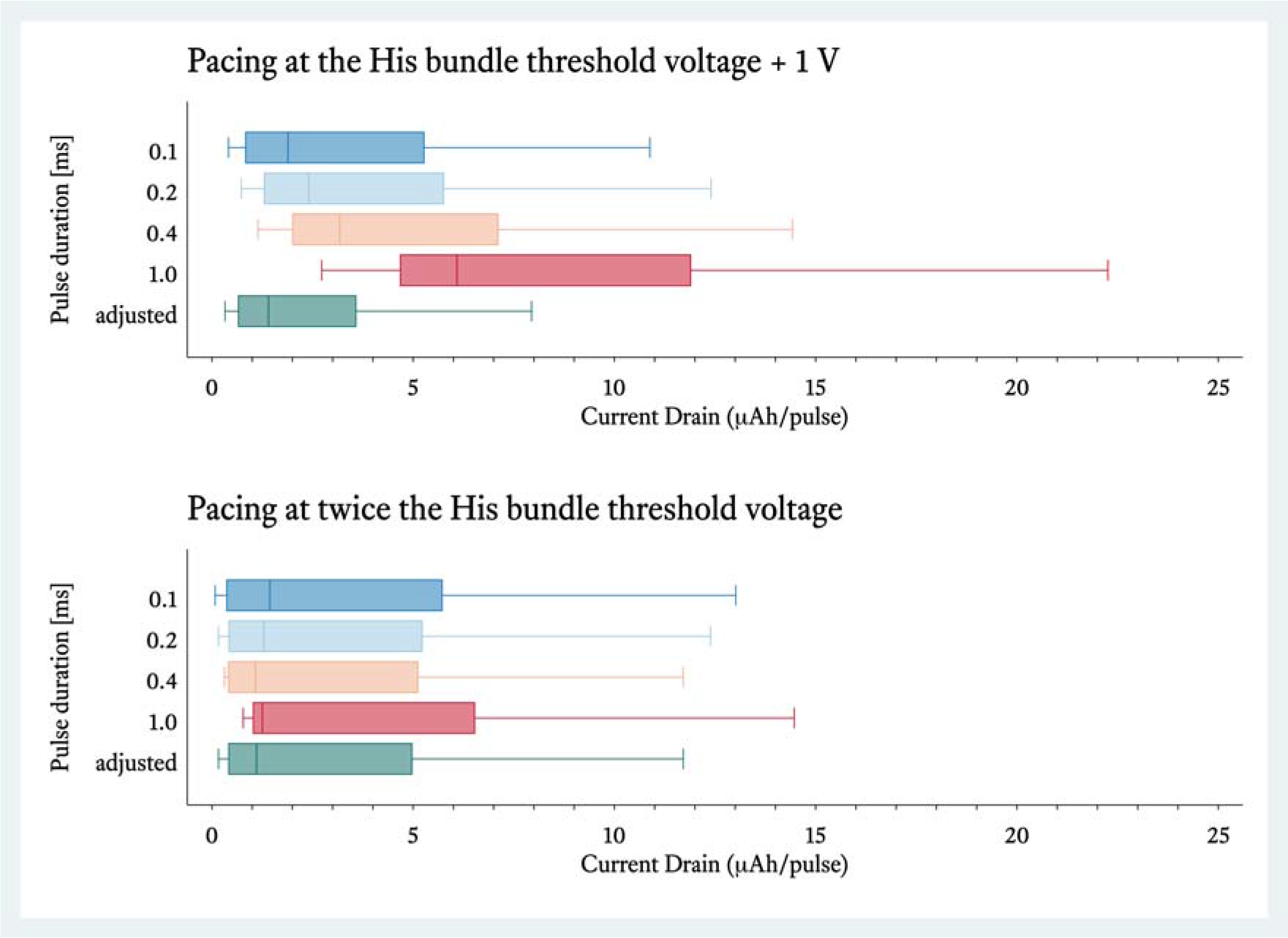
Battery current drain is lower with empirically programmed shorter pulse durations or chronaxie adjusted (see text) pulse duration. Box plot with median and interquartile range (25% - 75%).

## Discussion

This is the first study showing that HB chronaxie during permanent pacing with a screw-in lead is shorter that chronaxie of the adjacent RV myocardium. The current study also showed that this has potential practical implications as pacing with short pulse duration facilitates selective capture of the HB and results in less energy drain from the battery than pacing at 1 ms pulse width often used for permanent HB pacing.^17^

### Strength-duration curves and chronaxie values for RV and HB

Despite several studies reporting strength-duration curves for the RV myocardium, provided chronaxie values differ significantly and there is a scarcity of data with regard to the modern pacing leads. For example, while the often cited Coates and Thwaites paper, provided value of 0.24 ms for passive fixation, non-steroid eluting leads with cathode surface area of 6 mm^2^, Dhar et al. provided chronaxie value of 0.62 ms for more modern, steroid-eluting leads with active helix leads and cathode surface area of 5.7 mm^2^.^22, 23^ Both these studies investigated apical RV lead positions; data for septal lead position and active fixation leads with smaller cathode surface area are not available, to the best of our knowledge. Differences in reported chronaxie values might have various reasons, especially the type of lead used, cathode surface area, differences in electrophysiological properties in particular RV area, distance from the paced tissue and methodology of measurement. ^18, 24^ In the current study we have found that for the Medtronic 3830 lead at the HB capture position, the RV chronaxie is ms - close to the values obtained by Dhar et al., However, RV strength-duration curve when pacing from the 3830 lead strongly depended on the pacing lead position. In the obligatory non-selective HB capture group (i.e. when the RV threshold was always lower than the HB threshold), rheobase was 3 times lower than in patients with selective HB capture. This can be explained by the difference in distance of the pacing electrode (screwed-in helix) to the RV myocardium. In the obligatory non-selective HB capture patients it can be presumed that the pacing helix is directly within the RV working myocardium (parahisian position) as RV is preferentially depolarized. In selective HB group, this distance is likely bigger as the pacing helix is usually supravalvular,^25, 26^ within the atrioventricular septum, possibly directly penetrating the His bundle in some cases - resulting in limited contact with RV myocardium.

The HB chronaxie of 0.53 ms was found to be shorter than the adjacent local RV myocardial chronaxie. This might reflect the known relationship between chronaxie and velocity of conduction. Rapidly conducting tissues - like nerves, have shorter chronaxies (chronaxie of 0.05 ms) than slowly propagating tissues - like a denervated skeletal muscle (chronaxie of 9.5-30 ms). The velocity of conduction in HB is 3 – 5 times faster than in the working myocardium, it is, therefore, not surprising that the HB chronaxie was found to be shorter than the RV myocardium chronaxie. In our group, 25% of patients with selective HB pacing had chronaxie < 0.28 ms vs only < 5% of obligatory non-selective HB pacing. We believe that cases with the shortest chronaxie and lowest selective HB capture thresholds (lowest rheobase) represent possible intrahisian position — that most likely allow best approximation of true excitability of HB. The cases with longer HB chronaxie might represent suboptimal pacing lead positions with pacing helix at a larger distance from the HB.

Apart from our study, we are not aware of any other published data concerning HB chronaxie during permanent pacing. A study by Fozzard and Schoenberg concerning His bundle chronaxie assessed in the milieu of experimental ex vivo models reported chronaxie values of approximately 0.23 ms.^27^

### Selective vs non-selective HB capture

Although limited studies support the concept that non-selective HB pacing is hemodynamically equivalent to selective pacing,^28, 29^ we do not have concrete clinical evidence. Selective HB pacing is certainly closer to physiology and offers straightforward ECG diagnosis of presence/loss of HB capture and also allows to easily offset HV interval from atrioventricular delay during DDD pacing.^14^ Moreover, non-selective HB capture, especially in patients with prolonged His-Ventricle interval and or uncorrected bundle branch block occasionally results in a large degree of direct myocardial depolarization and very broad QRS complexes of >150 ms, and it is reasonable to suspect that such non-selective HB pacing may be hemodynamically inferior to selective HB pacing with narrower QRS complexes.

It was shown that stimulation of the vagus nerve with different pulse widths excites different nerve fibers within the nerve due to the differences in chronaxie between them.^30^ Similarly, in the pacing arena, differences in chronaxie were explored for selective pacing of the left ventricular myocardium and avoidance of phrenic nerve capture during cardiac resynchronization therapy. It was observed that the faster-conducting tissue (i.e. phrenic nerve) had shorter chronaxie than left ventricular myocardium (0.22 ms vs 0.47 ms). We believe that the phrenic nerve as a more distant structure has higher rheobase and pacing with the lowest possible amplitude avoids its capture during cardiac resynchronization therapy.^32^ In patients with permanent HB pacing the situation is reversed: the structure that we might desire to pace selectively (HB) has faster conduction velocity and shorter chronaxie than the structure that we might want to avoid capturing (RV). This is why selective capture is facilitated with pacing at shorter pulse widths at which the distance between HB and RV strength-duration curves is bigger since the steep portion of RV strength-duration curve starts later - see Figure 2. In other words, the zone of selective HB capture is widening with shortening of the pulse duration (Figure 2 and Figure 3). The current study shows that there is an increasing percentage of patients with selective HB pacing when shorter pulse widths are programmed (Figure 4). Of note, in our center we observed a higher percentage of patients with chronic selective HB pacing (59%) ^10^ than reported in studies by Zanon et al., Kronborg et al., and Sharma et al. with 28%, 12.5% and 45% of patients with selective pacing, respectively. These differences might be explained by differences in the customs of pulse width programming or studied population. It is our current practice to promote selective HB capture by empirical programming of pulse duration to 0.2 - 0.3 ms, unless this would result in pulse amplitude of > 2.8 V. However, if simultaneous backup RV capture is desired for clinical reasons (distal atrioventricular block) then long pulse duration (1.0 ms) should be preferred as it offers bigger safety margin for RV capture.

### Current drain during HB pacing

In the light of higher capture thresholds, lack of pacemakers dedicated for HB pacing with a larger battery and automatic capture algorithms capable to recognize HB capture, it is of paramount importance to optimize battery longevity with appropriate programming. Chronaxie values during HBP have a broader range than during conventional RV pacing. The default pacemaker values for pulse width: 0.4 – 0.5 ms or especially the 1.0 ms that is often used in HBP can be significantly far from the optimal (i.e. close to chronaxie). It has to be kept in mind that at the constant voltage, changing the pulse duration from 0.2 ms to 1.0 ms increases the battery drain 5 times. Using longer pulse widths might be deceptive as to the real current drain due to lower programmed output amplitude.

Our study shows that the lowest current drain can be achieved when the programming of pulse duration is guided by chronaxie, regardless of chosen safety margin (+ 1 V or twice voltage threshold).

### Limitation

Provided chronaxie values are for the Medtronic 3830 lead and might be different if a different lead is used for HB pacing.

Studied group consisted mainly of patients with narrow native QRS complexes who were not pacemaker dependent; in patients with complete infranodal heart block and/or left bundle branch block where recruitment of distal HP fibers is required, different HB chronaxie might be present. However, the few patients with infranodal block or left bundle branch block that were included in the selective HB group and had distal capture, did have short chronaxie. Although nonselective HBP may be preferred in this population for clinical reasons.

The impact of voltage multiplier on the current drain was not assessed. In patients with devices employing a voltage multiplier, programming a pulse width of 0.2 ms might result in voltage programmed to > 3 V, and, consequently, in higher current drain than when programming 1 ms pulse width and the amplitude < 3 V.

## Conclusions

Novel methods of cardiac pacing present new challenges and require thorough understanding of the basics of the new type of pacing. The current study shows that the permanent His bundle pacing is characterized by lower chronaxie and lower rheobase than RV pacing from the same lead. This might have practical implications, such that permanent His bundle pacing can be optimized with regard to obtaining selective HB capture and improving battery longevity by programming pulse durations according to estimated His bundle chronaxie.

